# G1899, an American ginseng extract, alleviates neuroinflammation and cognitive impairment in models of Alzheimer’s disease

**DOI:** 10.1101/2025.09.04.674314

**Authors:** Sin Ho Kweon, Jae-Hoon Lee, Byung-Cheol Han, Han Seok Ko

**Author notes:** Correspondence (H.S.K).

## Abstract

**Background:** Alzheimer’s disease (AD) is characterized by amyloid β (Aβ) accumulation, tau pathology, and chronic neuroinflammation, yet current therapeutic strategies provide only limited efficacy. Natural compounds with pleiotropic actions have emerged as potential adjunctive interventions. This study evaluated G1899, a standardized American ginseng (Panax quinquefolius) extract, for its effects on neuroinflammation, Aβ clearance, and cognitive function.

**Methods:** Neuronal cultures were exposed to glutamate or Aβ oligomers (Aβo) and pre-treated with G1899 to assess cell viability and excitotoxicity. Primary murine microglia were analyzed for validating expression levels of TMEM119, CD68, NLRP3 inflammasome, IL-1β, Caspase-1, and STAT3 phosphorylation. Behavioral testing was performed in scopolamine-injected mice with short term G1899 treatment and in 5xFAD transgenic mice following long-term G1899 administration at multiple doses. Amyloid burden, microglial recruitment, and plaque morphology were quantified by immunohistochemistry and high-resolution imaging. Human induced microglia (iMG) were examined for neuroinflammatory responses following Aβo exposure with or without G1899 treatment.

**Results:** G1899 significantly improved neuronal viability and reduced glutamate- and Aβo-induced toxicity. In microglia, G1899 upregulated TMEM119 and CD68, while suppressing NLRP3 inflammasome formation, proinflammatory cytokine expression, and STAT3 phosphorylation. G1899 rescued scopolamine-induced memory deficits and, in 5xFAD mice, reduced hippocampal and cortical Aβ burden, alleviated neuroinflammatory markers, and improved both spatial/fear learning and memory, with the most consistent efficacy observed at 300 mg/kg. Imaging revealed enhanced microglial recruitment to plaques and facilitated fragmentation of Aβ deposits. In iMG, G1899 elevated homeostatic and phagocytic markers while attenuating Aβo-induced NLRP3/STAT3-mediated neuroinflammatory signaling pathway.

**Conclusions:** G1899 confers multimodal neuroprotection by preserving neuronal survival, modulating microglial activity, and facilitating Aβ clearance. These findings highlight its potential as a safe and clinically translatable botanical intervention for AD.

## Background

Alzheimer’s disease (AD) is a progressive neurodegenerative disorder and the most common cause of dementia in the aging population. It is clinically characterized by a gradual decline in memory, executive function, orientation, increased extracellular amyloid β (Aβ) deposition, elevated accumulation of intracellular neurofibrillary tangles composed of hyperphosphorylated tau, chronic neuroinflammation, and synaptic degeneration [1]. These features contribute to structural disconnection and functional impairment in multiple brain regions. One of the primary contributors to the pathological cascade is the overproduction and aggregation of Aβ peptides, which result from sequential cleavage of amyloid precursor protein (APP) by β-secretase (BACE1) and γ-secretase. The catalytic core of the γ-secretase complex includes presenilin-1 (PSEN1), a protein that regulates the generation of Aβ42, a particularly aggregation-prone form of Aβ [2, 3]. Mutations in PSEN1 and abnormal upregulation of BACE1 have been implicated in familial and sporadic forms of AD, respectively [4, 5]. These alterations lead to excessive accumulation of Aβ in the brain, triggering a cascade of secondary pathological events including microglial activation, pro-inflammatory cytokine release, oxidative stress, and mitochondrial dysfunction [6, 7]. In addition to Aβ pathology, tau hyperphosphorylation destabilizes microtubules and disrupts axonal transport, contributing to synaptic and neuronal loss [8, 9]. Aβ also interferes with calcium homeostasis and synaptic vesicle cycling, impairing neurotransmitter release and long-term potentiation [10]. However, therapeutic efforts targeting BACE1 and γ-secretase have shown limited efficacy due to mechanism-related side effects, highlighting the need to understand upstream regulatory pathways of BACE1 and PSEN1 for more precise and tolerable interventions [11]. Given the limited clinical benefit of targeting Aβ or tau directly, interest has shifted toward natural compounds with multi-faceted therapeutic potential and lower risk of adverse effects. One such natural compound is Panax ginseng C.A. Meyer (ginseng) has attracted attention for its neuroprotective potential. Korean Red Ginseng (KRG), a heat-processed form of ginseng, contains multiple active ginsenosides, among which Rg3 has received particular attention [12, 13]. In preclinical studies, Rg3 reduced microglial activation [14, 15] and lowered pro-inflammatory cytokines such as TNF-α and IL-6 in the cortex and hippocampus [16]. KRG also restored synaptic proteins including PSD-95 and synaptophysin in the hippocampus [17] and improved memory performance in Y-maze and novel object recognition tests [18]. Furthermore, KRG has been tested as an adjuvant therapy in AD patients [19], and chronic administration of ginsenoside Rg1 improved spatial learning and memory in D-galactose-induced aging models [20]. Similarly, American ginseng (Panax quinquefolius L.), a medicinal herb indigenous to North America, has gained considerable attention for its potential to modulate cognitive performance and confer neuroprotective effects and emerged as a promising botanical candidate for cognitive enhancement. In double-blind, placebo-controlled trials involving healthy young and middle-aged adults, acute administration of standardized American ginseng extracts improved working memory, attention, and mental arithmetic performance within two to six hours post-ingestion [21, 22]. These effects have been attributed to modulation of cholinergic and glucose-regulatory pathways, supported by evidence from electrophysiological studies showing increased frontal cortical activation during cognitive tasks [23]. These outcomes were accompanied by normalization of hippocampal neurotransmitter levels, reduced oxidative stress, and restoration of mitochondrial function [24, 25]. Metabolomic analyses of cerebrospinal fluid from treated animals revealed reversal of AD-like metabolic signatures, suggesting that American ginseng affects systemic and central nervous system (CNS) homeostasis [26]. Studies in rats revealed no significant adverse effects on hepatic or renal function, and pharmacokinetic data suggests low systemic toxicity of its major ginsenosides such as Rb1 and Re [27, 28]. Collectively, these findings support American as a compelling therapeutic candidate with favorable safety profile for cognitive decline and neurodegenerative disorders. In this context, our study focuses on a novel American ginseng extract, G1899, to evaluate its potential as a neuroprotective agent in AD. Using both in vitro and in vivo models, such as human embryonic stem cell (H9)-derived differentiation of human microglia and 5xFAD transgenic mice, we examined the beneficial effects of G1899 on Aβ clearance, microglial activity, neuroinflammation, and cognitive deficits. Notably, G1899 has demonstrated significant improvements in cognitive performance in vivo models by alleviating neuroinflammation and microglial activation. Thus, we suggest for the first time that G1899 promotes Aβ degradation by enhancing microglial phagocytosis, which may reduce Aβ plaque burden and downstream neurodegeneration. Given its favorable safety profile and multi-modal actions, G1899 may represent a novel and clinically translatable therapeutic approach for AD patients, especially those unresponsive to current amyloid-targeting therapies.

## MATERIAL AND METHODS

### Animals

C57BL/6 mice were obtained from the Jackson Laboratory (ME, USA). 5xFAD mice, B6.Cg-Tg (APPSwFlLon,PSEN1*M146L*L286V)6799Vas/Mmjax, RRID:MMRRC_034848-JAX [29] were obtained from the Mutant Mouse Resource and Research Center (MMRRC) at The Jackson Laboratory, an NIH-funded strain repository, and the mouse was donated to the MMRRC by Robert Vassar, Ph.D., Northwestern University. All procedures involving animals were approved by and conformed to the guidelines of the Institutional Animal Care Committee of Johns Hopkins University (Protocol Number MO22M174). Mice were housed 5 per cage in a 12 h light/dark cycle with free access to water and food. All mice were acclimatized in the procedure room before starting any animal experiments. We have taken great measures to reduce the number of animals used in these studies, and all efforts were made to reduce animals suffering from pain and discomfort.

### Primary murine cortical neuron and microglia culture

Cortical neurons were prepared from embryonic day 15.5 of pregnant CD-1 mice (Charles River, Wilmington, MA). Dissected neurons were plated onto dishes pre-coated with poly-L-lysine (PLL, Sigma Aldrich) in Borate buffer (50mM Boric acid in ddH2O, pH 8.5). Primary cortical neurons were cultured in complete medium (NM0) consisting of Neurobasal media (Invitrogen), containing B27 supplement, 1x GlutaMAX and 1x penicillin/streptomycin (Gibco) on tissue culture well plates coated with 100 μg/ml poly-D-lysine (Sigma-Aldrich). The NM0 was freshly added to cell culture every three days. All procedures involving mice were approved by and conformed to the guidelines of the Johns Hopkins University Animal Care and Use Committee. Whole brains from mouse pups at postnatal day 1 (P1) were obtained. After removal of the meninges, the brains were washed in DMEM/F12 (Gibco) supplemented with 10% heat-inactivated FBS, 50 U/ml penicillin, 50 μg/ml streptomycin, 2 mM L-glutamine, 100 μM non-essential amino acids and 2 mM sodium pyruvate (as a complete medium) three times. The brains were transferred to 0.25% trypsin-EDTA followed by 10 min of gentle agitation. DMEM/F12 complete medium was used to stop the trypsinization. The brains were washed three times in this medium again. A single-cell suspension was obtained by trituration. Cell debris and aggregates were removed by passing the single-cell suspension through a 100-μm nylon mesh. The final single-cell suspension thus achieved was cultured in T75 flasks per brain for 13 days. The mixed glial cell population was separated into microglia-rich fractions using the EasySep Mouse CD11b Positive Selection Kit (STEMCELL Technology #18970) in accordance with the manufacturer’s instructions. The magnetically separated fraction containing microglia from the non-binding fraction was cultured. After 24 h, microglia were kept in FBS-free DMEM/F12 supplemented with 50 U/ml penicillin, 50 μg/ml streptomycin, 2 mM L-glutamine, 100 μM non-essential amino acids, and 2 mM sodium pyruvate. The amyloid β was purchased from rPeptide (Beta-Amyloid (1-42), HFIP, 1mg, #A-1163-2) and was reconstituted with 100µl DMSO followed by 900µl of DPBS to make amyloid β oligomer by using microtube shaking incubator at 37°C for 1h at 1000rpm.

### *In vitro* immunofluorescence analysis

Primary microglia (3.0 × 10^5^ cells/well in 24-well plates) were seeded on coverslips coated with Poly-L-lysine (PLL) in Borate buffer for 24 h. Primary microglia was treated with 5μM of Aβo with or without 12 hours pretreated G1899. After 24hours of Aβo treatment, cells were washed with PBS and fixed with 4% paraformaldehyde for 10 min at RT, permeabilized with 0.1% Triton X-100 in PBS for 20 min and washed with DPBS three times. Then, cells were blocked with 5% normal goat serum (Jackson ImmunoResearch) in 0.1% Tween 20 in PBS (PBST) for 30 min and incubated with anti-Iba1 (1:500, Wako, #019-19741), anti-TMEM119 (1:500, Biolegend, #853302) at 4 °C overnight. Cells were then washed with PBST for three times followed by incubation in a mixture of Alexa Fluor 488 (1:1000, ThermoFisher, #A11001) and Alexa Fluor 568 (1:1000, ThermoFisher, # A11011) secondary antibodies for 1hr at room temperature and then washed three times with PBST. Coverslips were mounted on a glass slide in a mounting medium with DAPI (VECTASHIELD, Vector Laboratories, CA, USA). The fluorescence images were acquired via a Zeiss confocal microscope (LSM 880/900, Carl Zeiss, CA, USA).

### Immunohistochemistry and immunofluorescence

Mice were fixed with 4% paraformaldehyde/PBS (pH 7.4) after receiving an ice-cold PBS perfusion. After being collected, the brains were post-fixed for 16 h in 4% paraformaldehyde and followed by cryoprotected in a solution of 30% sucrose/PBS (pH 7.4) for 3 days. An optimal cutting temperature (OCT) buffer was used to freeze the brains before cutting 30 µm serial coronal pieces with a cryostat (LEIKA CM3050S). Free-floating 30 µm sections were incubated with an antibody against Aβ (4G8) (1:1,000, Biolegend, #800708) followed by incubation with a biotin-conjugated anti-rabbit or anti-mouse antibody (1:250, Vectastain Elite ABC kit, Vector laboratories). Sections were counterstained with Nissl (0.09% thionin) after being developed using SigmaFast DAB Peroxidase Substrate (Sigma-Aldrich) followed by usage of a computer-assisted image analysis system composed of an Axiophot photomicroscope (Carl Zeiss Vision) fitted with a computer controlled motorized stage (Ludl Electronics), a Hitachi HV C20 camera, and Stereo Investigator software (MicroBright-Field). For immunofluorescence, 4% Paraformaldehyde/PBS (pH 7.4)-fixed 30 µm coronal brain sections in SNc, hippocampus, and amygdala regions were blocked with 10% goat serum (Jackson Immunoresearch)/PBS plus 0.1% Triton X-100 and incubated with antibodies to 4G8 (1:1,000, Biolegend, #800708), NeuN (1:1,000, Millipore, #MAB377), and Iba1 (1:1,000, Wako, #019-19741) overnight at 4°C. Following PBS washes, floating brain sections were incubated with 0.1% Triton X-100 and 5% goat serum in PBS with a mixture of Alexa Fluor 488 (1:1,000, ThermoFisher, #A11001) and Alexa Fluor 568 (1:1,000, ThermoFisher, #A11011) secondary antibodies for 1 h at room temperature and then washed three times with PBST. After mounting the coverslips with DAPI mounting medium (VECTASHIELD HardSet Antifade Mounting Medium with DAPI, Vector laboratories), the fluorescence images were captured using a Zeiss confocal microscope (LSM 880/900). The Zeiss Zen software was used to process each image. ImageJ analysis was used to calculate the size of the chosen region inside the threshold’s signal intensity range.

### Immunoblot analysis

Mouse primary microglia and hippocampus region of mouse brain from control and 5xFAD mice with or without G1899 at concentration of 300mg/kg were lysed in RIPA lysis buffer (25 mM Tris•HCl pH 7.6, 150 mM NaCl, 1%NP-40, 1% sodium deoxycholate, 0.1% SDS) containing protease inhibitor cocktail (Complete^TM^ EDTA-free protease inhibitor cocktail, Roche) and phosphatase inhibitor (PhosSTOP^TM^, Roche) followed by centrifugation at 12000rpm for 10 min in 4 °C to collect supernatants. Protein concentration was measured using BCA Kit (Pierce, IL, USA). Equal amounts of proteins (10-20μg) prepared and then mixed with 4X Laemmli buffer (Bio-Rad) to resolve the sample on 8-16% Tris-Glycine or 4-12% Bis-Tris gradient gels followed by transfer to nitrocellulose membranes. These membranes were blocked with blocking solution (Tris-buffered saline with 3% or 5% non-fat dry milk respectively based on antibody specificity with 0.05% Tween-20) for 1 h followed by incubation at 4 °C overnight with primary antibodies; anti-Oct-4A (1:1000, Cell signaling Technology (CST), #2840T), anti-Nanog (1:1000, CST, #4903T), anti-CD34 (1:1000, CST, #26233S), anti-iNOS (1:1000, Novus Biologicals, #NB300- 605), anti-NLRP3 (1:1000, CST, #13158S), anti-TMEM119 (1:1000, Biolegend, #853302), anti-human CD68 (1:500, Biolegend, #375602), anti-mouse CD68 (1:500. Biolegend, #137001), anti- CD68 (1:1000, CST, #9778S), anti Caspase1 (1:500, Adipogen Life Sciences, AG-20B-0044- C100), anti-ASC (1:1000, Adipogen Life Sciences, AG-25B-0006), anti-Iba1 (1:1000, Wako, #019- 19741), anti-pY705 STAT3 (1:1000, CST, #9145S), anti-pS727 STAT3 (1:1000, CST, #9134S), anti-total STAT3 (1:1000, CST, #9139S), anti-Presenilin1 (1:1000, CST, #3622S), anti-APP (1:1000, MilliporeSigma, #MAB348), anti-Syntaxin 1A (1:1000, Biolegend, #827001), anti-IL-1β (1:1000, CST, #12242S), anti-Aβ (4G8) (1:1,000, Biolegend, #800708), anti-Aβ (6E10) (1:1,000, Biolegend, #803014) or anti-β-actin (1:5000, Sigma, #A3854-200UL) followed by HRP-conjugated rabbit (#32260) or mouse (#32230) secondary antibodies (1:5000, Thermo) for 1 hr at room temperature. Chemiluminescence signals from immunoblot were visualized by ImageQuant LAS4000 mini (Cytiva) or Amersham Image Quant 800 (Cytiva).

### Scopolamine induced cognitive decline

To evaluate the beneficial effect of G1899 on elevating cognitive ability from memory impairment induced by intraperitoneal injection (i.p.) of Scopolamine (2mg/kg), a selective muscarinic acetylcholine receptor antagonist, widely used for mimicking learning and memory deficit. The animals were divided into control groups receiving distilled water, Scopolamine (SCP) receiving group, G1899 receiving group at 300mg/kg via oral administration, and a positive control group receiving Donepezil (5mg/kg) with oral injection. After 4 weeks of oral administration period (5days per week), SCP, G1899, Donepezil groups were co-injected with SCP at 2mg/kg during the first and second day of training followed by actual testing on the third day. For validating efficacy of G1899 against scopolamine (SCP, 1mg/kg) induced memory impairment test, 3 mo. aged mice were used and treated with vehicle, 30, 100, 300mg/kg of G1899 daily prior to fear conditioning test. During the training and actual test session, mice were intraperitoneally injected with 1mg/kg SCP before admitting to fear conditioning test chamber.

### Fear conditioning (FC)

We performed fear conditioning test (Med Associates, Inc., VT, USA) for validating cued learning and memory. A conditioned stimulus (tone), an aversive unconditioned stimulus (shock), new olfactory cue (acetic acid), floor texture (grid to acryl plate), visual cue (light on/off) were used. On the first day, the mice were placed in chamber of context A for 10 min to habituate. On the second day, mice received four tone-shock pairings, and the shock (0.5 mA, 3 sec) was applied after the end of the tone (80 dB, 5 kHz, 15 sec). On the 4th day, mice were either placed in context A for 5 min without tone and shock for contextual FC or context B with different textures, olfactory and visual cues for cued FC with tone only. In cued FC, after tone alarming, the freezing episode and total freezing time was measured and validated as a cued FC. Freezing behavior is defined as the complete lack of motion for a minimum of 1 second.

### Morris Water Maze (MWM)

The Morris water maze (MWM) is a test of spatial learning and memory for rodents in an open round swimming arena that is 150 cm in diameter and 50 cm high. and it has four separate inner cues right above the water surface. The platform was submerged 1∼2 cm below the water surface so that it is invisible at water level. The circular pool was filled with water and a nontoxic water-soluble white dye (20 ± 1°C). The swimming pool was divided into four equal-sized quadrants. Within one of the quadrants, a platform (9 cm in diameter and 15 cm high) was positioned in the middle of target quadrant. Using a video tracking system (ANY-Maze), movement of mouse by swimming to the platform region is recorded by automatic mouse tracking system. The mice were subsequently subjected to three trial sessions (3 different starting points from certain quadrant except for target quadrant) per day during consecutive four days, then the escape latencies were recorded. For each trial session and each mouse, this parameter was averaged. The mouse was given permission to stay on the platform for 10 s after finding it. If the mouse failed to locate the platform in 60 s, it was put there for 10 s before being put back in its cage by the experimenter. On day 6, mice were given the probe trial test that involved removing the platform from the water.

### Culture and differentiation of hESCs into microglia

All experiments involving human embryonic stem cells (hESCs) were conducted in compliance with protocols approved by the Johns Hopkins Medicine Institutional Review Board and the Institutional Stem Cell Research Oversight (ISCRO) Committee at Johns Hopkins University (Approval No. ISCRO00000620). Human H9 embryonic stem cells (WA09, NIHhESC-10–0062) were obtained from WiCell and were plated on Geltrex and maintained in mTeSR1 or mTeSR1 plus medium throughout the cell expansion. According to manufacturer’s protocol for STEMdiff^TM^ hematopoietic kit (STEMCELL Technology #05310), human ESC colonies were harvested and seeded as small aggregates followed by the replacement of mTeSR1 medium with Hematopoietic Medium A to induce the cells toward a mesoderm-like state. On Day 3, change to Medium B and perform half-medium changes on Days 5, 7, and 10 to promote further differentiation into hematopoietic cells. And by Day 12, large numbers of hematopoietic progenitor cells can be harvested from the culture supernatant. On the day of microglia differentiation STEMdiff^TM^ microglia differentiation (Cat #100-0019, up to 24 days) and maturation (Cat #100-0020, up to 12 days) kit were used based on the manufacturer’s instruction.

### Statistics

All data from at least 3 separate experiments were given as mean ± standard error of mean (SEM). Differences between two groups were analyzed using one-way analysis of variance (ANOVA) with Tukey post hoc test. Two-way ANOVA performed to test for significant effects of genotype and G1899 treatment. The Tukey post hoc test was used for post-hoc multiple comparisons. The values of P represent the strength of the evidence supporting differences between groups for descriptive purposes. Consequently, these comparisons were not hypothesis testing and no correction was made for multiple hypothesis testing. All analyses were performed using GraphPad Prism 10 (GraphPad Software Inc.).

### Neuroprotective effect of G1899 by modulating amyloid beta oligomer (Aβo)-mediated inflammation via NLRP3/STAT3 pathway

To assess the neuroprotective effects of G1899, primary murine neurons were treated with glutamate (10µM) or amyloid beta oligomer (Aβo, 2.5µM) in the presence or absence of G1899. Firstly, cell viability was conducted using MTT assay. In the group pre-treated with G1899, cell death was reduced while promoting cell viability **(Fig. S1a, b).** Additionally, we examined the effect of G1899 on Aβo-induced neuroinflammation using primary murine microglia **(Fig. 1a)**, as Aβo are known to play a pivotal role in neuroinflammation of the central nervous system (CNS). To this end, primary microglia were treated with varying concentrations of G1899 to examine its impact on microglial activation and phagocytic function. Western blot (WB) analysis was conducted to assess the protein levels of transmembrane protein 119 (TMEM119) and cluster of differentiation 68 (CD68) [30], markers for beneficial activation and phagocytosis in microglia respectively. G1899 treatment significantly increased both proteins in a dose dependent manner **(Fig. 1b-d)**. Furthermore, we examined whether G1899 alleviates microglial activation under in vitro conditions mimicking AD, where primary microglia were exposed to Aβo. Aβo-treated microglia showed a substantial increase in NLRP3 and related neuroinflammatory signaling markers, including caspase 1 and apoptosis-related speck-like protein containing a caspase recruitment domain (ASC) **(Fig. 1e)** [31, 32]. In contrast, the treatment of G1899 significantly decreased the expression of NLRP3 as well as its downstream markers **(Fig 1f-h)**. Specifically, Aβo-treated microglia showed increased phosphorylation of pY705 STAT3 and pS727 STAT3, which was significantly reduced by G1899 treatment **(Fig 1i-k)**. The morphology of Aβo-treated microglia showed a typical amoeboid microglial shape commonly observed in various CNS-related disease models as assessed by immunocytochemistry. In contrast. These alterations were rescued by G1899 treatment **(Fig 1l, m)**. Furthermore, Aβo-treated microglia showed aberrant microglial activation, with increased in inducible nitric oxide synthase (iNOS), another specific microglia activation marker [33], and reduced TMEM119. These alterations were restored by G1899, as assessed by WB analysis **(Fig 1n-p)**.

**Fig 1.**
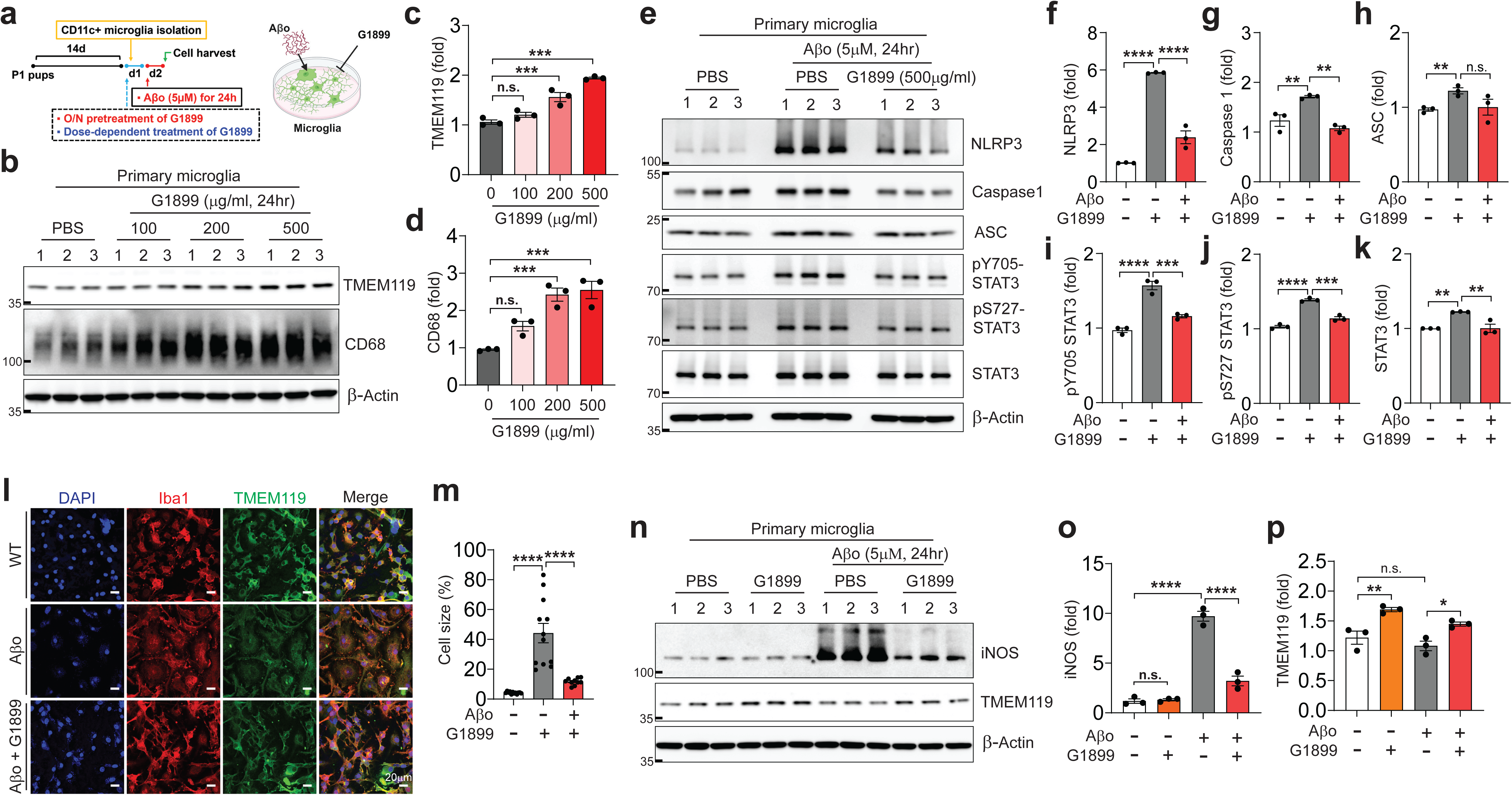
G1899 promotes microglial homeostasis and attenuates detrimental microglial activation induced by amyloid β oligomer (Aβo). **a** Schematic illustration (created with BioRender.com) of the treatment time window for G1899 and Aβo in mouse primary microglia. **b** Representative immunoblot images showing dose-dependent upregulation of anti-TMEM119 (homeostatic marker) and anti-CD68 (phagocytic marker) in G1899-treated microglia. Quantification of **c** TMEM119 and **d** CD68 normalized to β-actin. **e** Effects of G1899 on NLRP3 inflammasome formation and STAT3 signaling, with quantification of **f** NLRP3, **g** Caspase-1, **h** ASC, **i** pY705 STAT3, **j** pS727 STAT3, and **k** total STAT3 normalized to β-actin. **l** Immunofluorescence analysis of morphological changes using anti-TMEM119 and anti-Iba1 staining. Scale bar, 20 μm. **m** Quantification of cellular area following Aβo exposure, demonstrating restoration of microglial morphology by G1899. **n** Representative immunoblot images of anti-iNOS (activation marker), anti-TMEM119, and β-actin with quantification of **o** iNOS and **p** TMEM119 normalized to β-actin. Error bars indicate mean ± SEM. Statistical significance is determined by one-way ANOVA (c, d, f, g, i-p) or two-way ANOVA (h) with Tukey’s multiple comparisons test. *p < 0.05, **p < 0.01, ***p < 0.001, ****p < 0.0001; n.s., not significant.

### Beneficial effect of G1899 on restoration of cognitive impairment induced by scopolamine in vivo

Scopolamine (SCP, 2mg/kg) is widely used to induce acute memory impairment by disrupting the cholinergic system through muscarinic acetylcholine receptors antagonism, thereby mimicking amnesia and AD like pathology [34, 35]. To evaluate the effect of G1899 against acute memory loss, we employed a SCP-induced cognitive impairment and neurodegeneration model in mice. G1899 was administrated for 4 weeks before SCP treatment, and cognitive function was assessed using the fear conditioning (FC) test **(Fig. S2)**. Donepezil (5mg/kg, PO) was used as a positive reference to assess its rescue effect. SCP was injected right before each session of FC experiments, including habituation, training, and actual testing. Freezing behavior was assessed to determine fear memory retention. Mice treated with SCP (2mg/kg) alone showed a significant reduction in freezing behavior, whereas mice treated with G1899 (300mg/kg), or donepezil along with SCP showed restored percent freezing time **(Fig. S2)**. Additionally, to determine whether G1899 could rescue acute cognitive deficit in a dose-dependent manner, we administered three concentrations of G1899 (30, 100, and 300 mg/kg) along with a slightly lower dose of SCP (1 mg/kg, i.p.). A new set of FC experiments was performed using 2- to 3-month-old mice as schematically outlined in **Fig. 2a**. Although a clear dose-dependent trend was not observed, a partial rescue pattern was observed in the contextual fear conditioning test, which did not reach statistical significance **(Fig 2b)**. In the cued fear conditioning test, no clear dose-dependent improvement was observed, but cognitive rescue was significantly observed at the high dose of 300 mg/kg of G1899 **(Fig. 2c)**. These results suggest that G1899 exerts a partial restorative effect in the scopolamine-induced acute cognitive impairment and promotes neuronal resilience as a preventive intervention.

**Fig 2.**
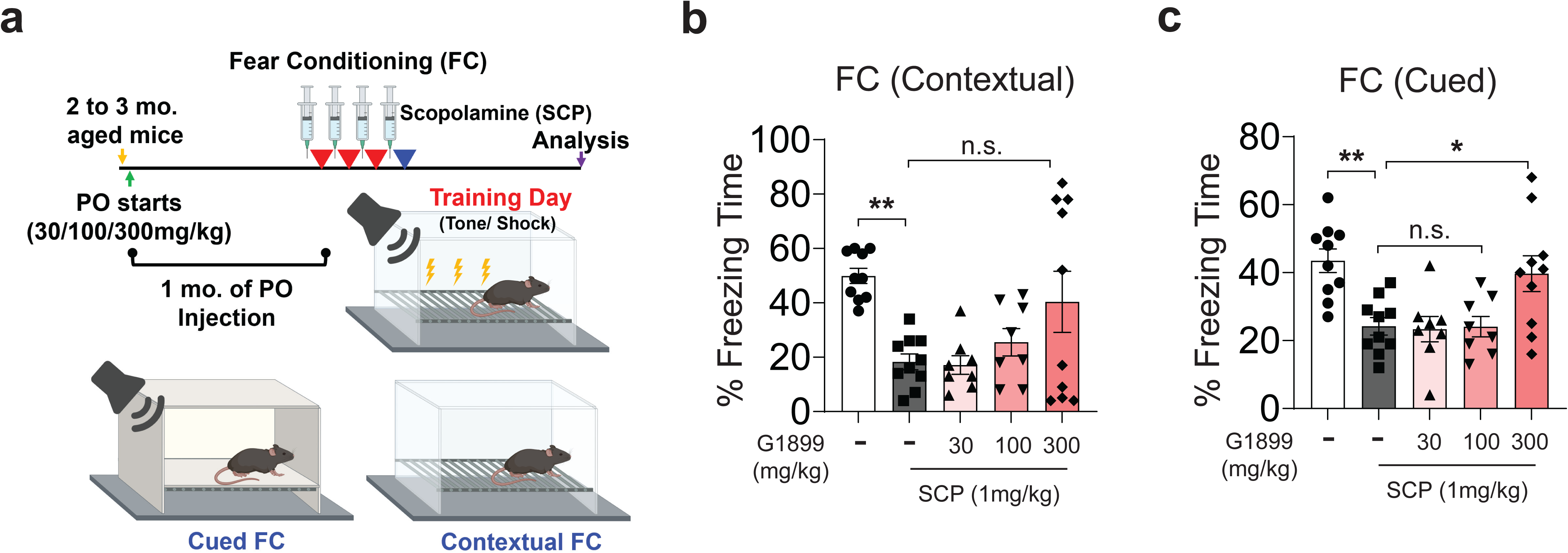
Neuroprotective effect of G1899 in a scopolamine-induced acute memory deficit model. **a** Schematic illustration of cued and contextual fear conditioning test (created with BioRender.com). Quantification of **b** contextual and **c** cued fear conditioning tests. Error bars indicate mean ± SEM. Statistical significance is determined by one-way ANOVA followed by Tukey’s multiple comparisons test. *p < 0.05, **p < 0.01; n.s., not significant.

### Beneficial effect of G1899 on restoration of cognitive impairment in 5xFAD mice

In addition to the SCP-induced mouse model, we next examined whether long term administration of G1899 could prevent AD pathology utilizing 5xFAD/+ mice, a well-established mouse model of AD. G1899 was orally administered to 2 to 3-month-old mice for six months, after which cognitive behavioral tests were performed and brain tissues were collected for biochemical assays from the respective groups of mice **(Fig. 3a)**. The experimental groups were categorized as follows: 1) WT vehicle, 2) WT 300 mg/kg, 3) 5xFAD/+ vehicle, 4) 5xFAD/+ 30mg/kg 5) 5xFAD/+ 100mg/kg 6) 5xFAD/+ 300mg/kg. To evaluate the potential for dose-dependent recovery, we first conducted a fear conditioning test. In the cued FC, G1899-treated 5xFAD/+ groups showed a significant increase at all dose levels compared to the vehicle-treated 5xFAD/+ group **(Fig. 3b)**. In the contextual FC, a significant rescue of cognitive performance was specifically observed at the highest dose of 300 mg/kg **(Fig. 3c)**. Furthermore, in the Morris water maze test, the representative tracking plots from the final training day demonstrated that the 5xFAD/+ mice treated with G1899 accurately remembered the platform’s location, finding it quickly and efficiently compared to vehicle-treated 5xFAD/+ **(Fig. 3d)**. On the day of the probe trial, the 5xFAD/+ mice treated with G1899 also spent a longer duration in the target quadrant, as confirmed via heat map **(Fig. 3e)**. Group-wise analysis revealed that the high-dose group showed a particularly faster and more efficient tendency to visit and remain in the target quadrant. To further examine whether the mice could precisely locate the probe, we analyzed additional regions such as 1) target quadrant, 2) an actual platform, and 3) extended platform. The high-dose group of 5xFAD/+ mice consistently remembered the designated platform area, showing higher number of visits **(Fig. 3f-h)** and longer stay **(Fig. 3i-k)** in both the actual and extended platform regions, demonstrating restored spatial learning and memory. These behavioral experiments confirm that G1899 administration effectively recover cognitive impairment in 5xFAD/+ mice compared to the vehicle-treated group.

**Fig 3.**
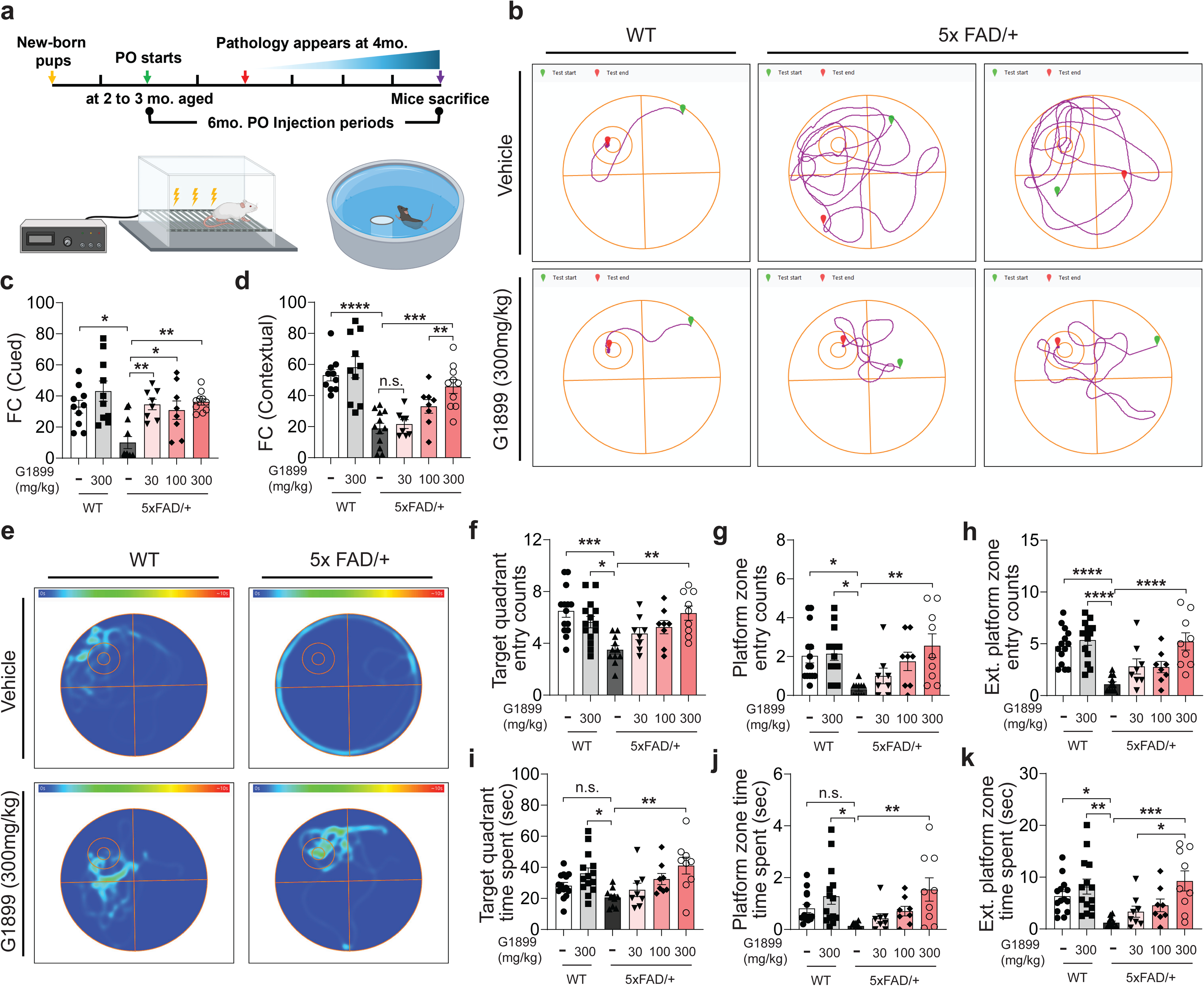
Cognitive improvement by G1899 in 9-month-old 5xFAD/+ mice. **a** Schematic illustration of the oral gavage (PO) dosing schedule of G1899 at different doses (30, 100, 300 mg/kg), administered five times per week for six months followed by behavior assessments. The images were created with BioRender.com. **b** Cued and **c** contextual fear conditioning tests assessing the effects of G1899 on cognitive improvement in 5xFAD/+ mice. **d** Morris water maze test used to evaluate the rescue effect of G1899. Representative tracking plots of swimming paths to the platform at the end of training (day 4). **e** Heat maps from each group during the probe trial (day 5). Quantification of the number of entries into the **f** target zone, **g** platform zone, and **h** extended platform zone, as well as time spent in the **i** target zone, **j** platform zone, and **k** extended platform zone. Error bars indicate mean ± SEM. Statistical significance is determined by one-way ANOVA followed by Tukey’s multiple comparisons test. *p < 0.05, **p < 0.01, ***p < 0.001, ****p < 0.0001; n.s., not significant.

### Reduction of amyloid β (Aβ) plaque in brain regions governing cognition by administration of G1899 in 5xFAD mice

To examine the beneficial effects of G1899 on Aβ plaque clearance, we collected the hippocampus (HPC), prefrontal cortex (PFC), and entorhinal cortex (EC) from both control and experimental groups, where considered as critical disease lesions associated with Aβ plaque formation in the 5xFAD mouse model of AD. Elevated levels of Aβ in the 5xFAD mice were significantly reduced in three different regions of brain from G1899-treated groups **(Fig. 4a-f)**. Although a clear dose-dependent reduction was not observed, the results demonstrated a significant decrease in Aβ levels in the high-dose group (300 mg/kg), along with meaningful reductions at lower dose levels. In addition, analysis of coronal brain sections encompassing the dorsal subiculum of the hippocampus revealed that immunohistochemistry targeting Aβ plaque (4G8) showed a widespread increase in 5xFAD/+ mice compared to wild-type controls, whereas the G1899-administered group exhibited a general reduction in Aβ deposition **(Fig. S3)**. Consistently, immunofluorescence imaging demonstrated a marked decrease in intensity and size of Aβ in the vicinity of the dorsal subiculum when co-stained with NeuN and Aβ **(Fig. 4g-i)**. These findings suggest the potential of G1899 as a preventive therapeutic agent against Aβ accumulation. We further examined Alzheimer’s disease (AD)-associated pathological markers, including amyloid precursor protein (APP), presenilin-1 (PSEN1), and Iba1, to evaluate microglia-driven neuroinflammatory processes **(Fig. S4)**. Our analyses demonstrated that G1899 mitigated not only conventional AD pathological markers but also those indicative of microglial activation and neuroinflammation. Moreover, the rescue effects previously established in primary microglial cultures were recapitulated in vivo, underscoring the capacity of G1899 to attenuate neuroinflammatory signaling and highlighting its potential as a modulator of AD-related neuroimmune pathology.

**Fig 4.**
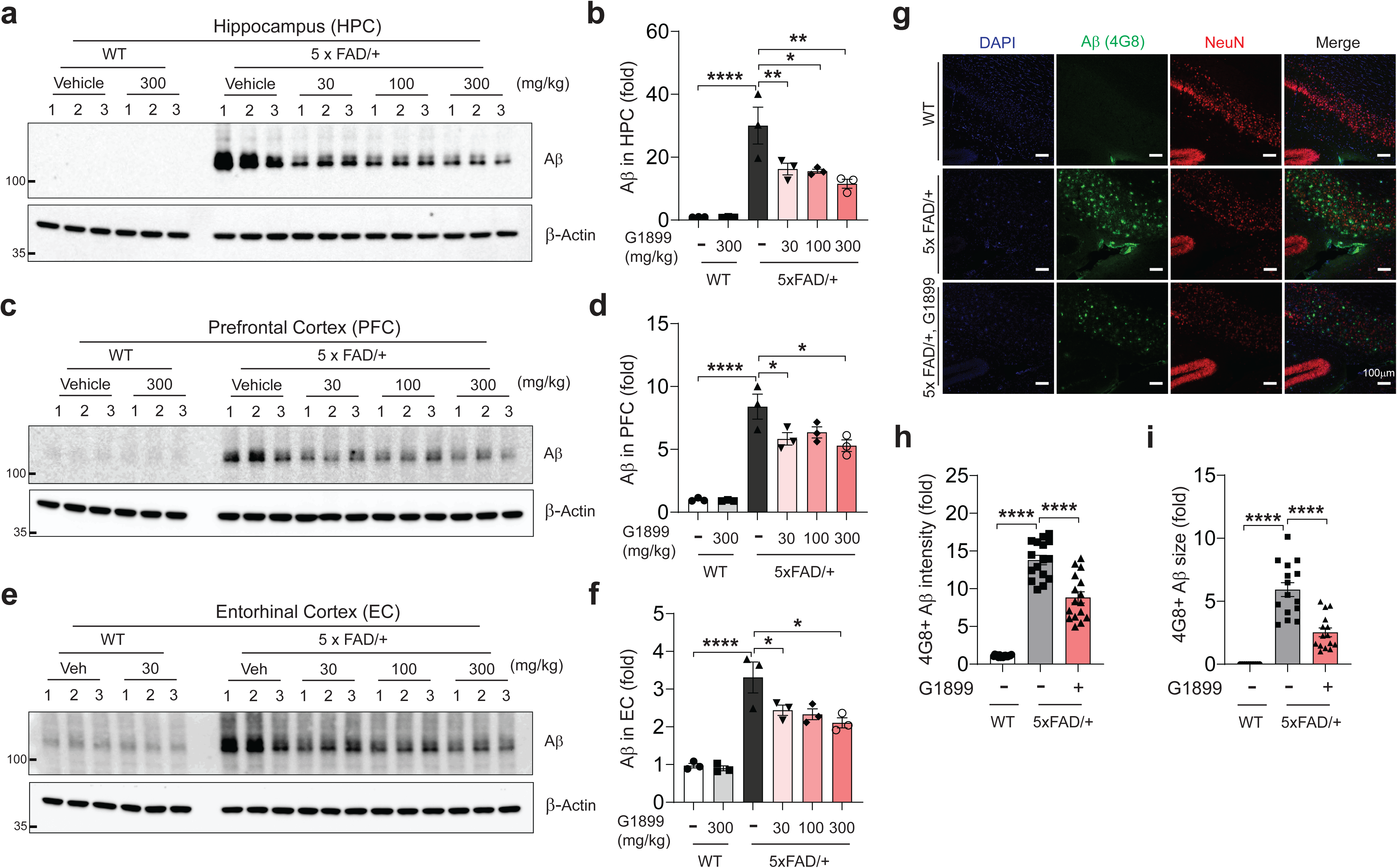
Reduction of Aβ accumulation in hippocampus (HPC), prefrontal (PFC), and entorhinal cortices (EC) by G1899. Representative immunoblot images from **a** HPC, **c** PFC, and **e** EC. Quantification of Aβ levels in **b** HPC, **d** PFC, and **f** EC in WT vehicle, WT with 300 mg/kg G1899, 5xFAD/+ vehicle, and 5xFAD/+ treated with 30, 100, or 300 mg/kg G1899. **g** Immunofluorescence assay detecting Aβ (4G8 clone) in the dorsal subiculum of coronal brain sections from WT vehicle, 5xFAD/+ vehicle, and 5xFAD/+ treated with 300 mg/kg G1899. Scale bar, 100 μm Quantification of **h** immunofluorescence intensity and **i** size of 4G8+ Aβ plaques. Statistical significance is determined by one-way ANOVA followed by Tukey’s multiple comparisons test. *p < 0.05, **p < 0.01, ***p < 0.001, ****p < 0.0001; n.s., not significant.

### Reduction of NLRP3 inflammasome via phospho-STAT3 pathway by administration of G1899 in 5xFAD mice

Based on the evidence suggesting G1899 as beneficial modulator for neuroinflammation, we aimed to observe the patterns of neuroinflammatory signaling pathways in the disease model and determine whether the anti-inflammatory signaling pathway was activated in the G1899-treated group, thereby playing a role in disease suppression. Upon examining the neuroinflammatory changes in the hippocampus, we found that the amount of Aβ plaque in the 5xFAD/+ group treated with G1899 was significantly reduced by approximately 20% to 30% **(Fig. 5a, b)**. Moreover, in relation to neuroinflammation, IL-1β **(Fig. 5c)** and caspase 1 **(Fig. 5d)** signaling, which were significantly elevated in 9-month-old 5xFAD/+ mice, whereas decreased in the G1899-treated group. Similarly, validation of involvement of STAT3 by measuring phosphorylated Y705 STAT3, which was also increased in the 5xFAD/+ mice but eventually decreased in the G1899-treated group **(Fig. 5e)** as well as reactive microglial activation determined by Iba1 was restored in G1899-administered group **(Fig. 5f)**. Notably, neurodegeneration validated by syntaxin 1A [36] **(Fig. 5g)**, presynaptic maker, demonstrated that neuronal survival was partially restored in G1899-treated group. To assess the generation of Iba1-positive reactive microglia upon Aβ plaque exposure, the fluorescence intensity of Iba1 was measured. It was elevated in 5xFAD/+ mice, whereas it was downregulated in G1899 treated group (**Fig. 5h, j**). To determine whether G1899 enhances microglia-mediated clearance of Aβ, z-stack images were acquired at 0.2 μm intervals across a depth of 5–10 μm based on Aβ signals, followed by projection into single composite images **(Fig. 5i)**. Quantification revealed a higher number of microglia localized in proximity to Aβ plaques in G1899-treated 5x FAD/+ groups compared with untreated group. In addition, compact circular Aβ plaques were fragmented into smaller structures, accompanied by significant reductions in plaque size and signal intensity, indicating that G1899 promotes microglial migration and recruitment to sites of Aβ deposition **(Fig. 5k)**. In conclusion, we speculated that G1899- treated group recovered phagocytic activity in comparison with non-treated group to facilitate Aβ plaque degradation.

**Fig 5.**
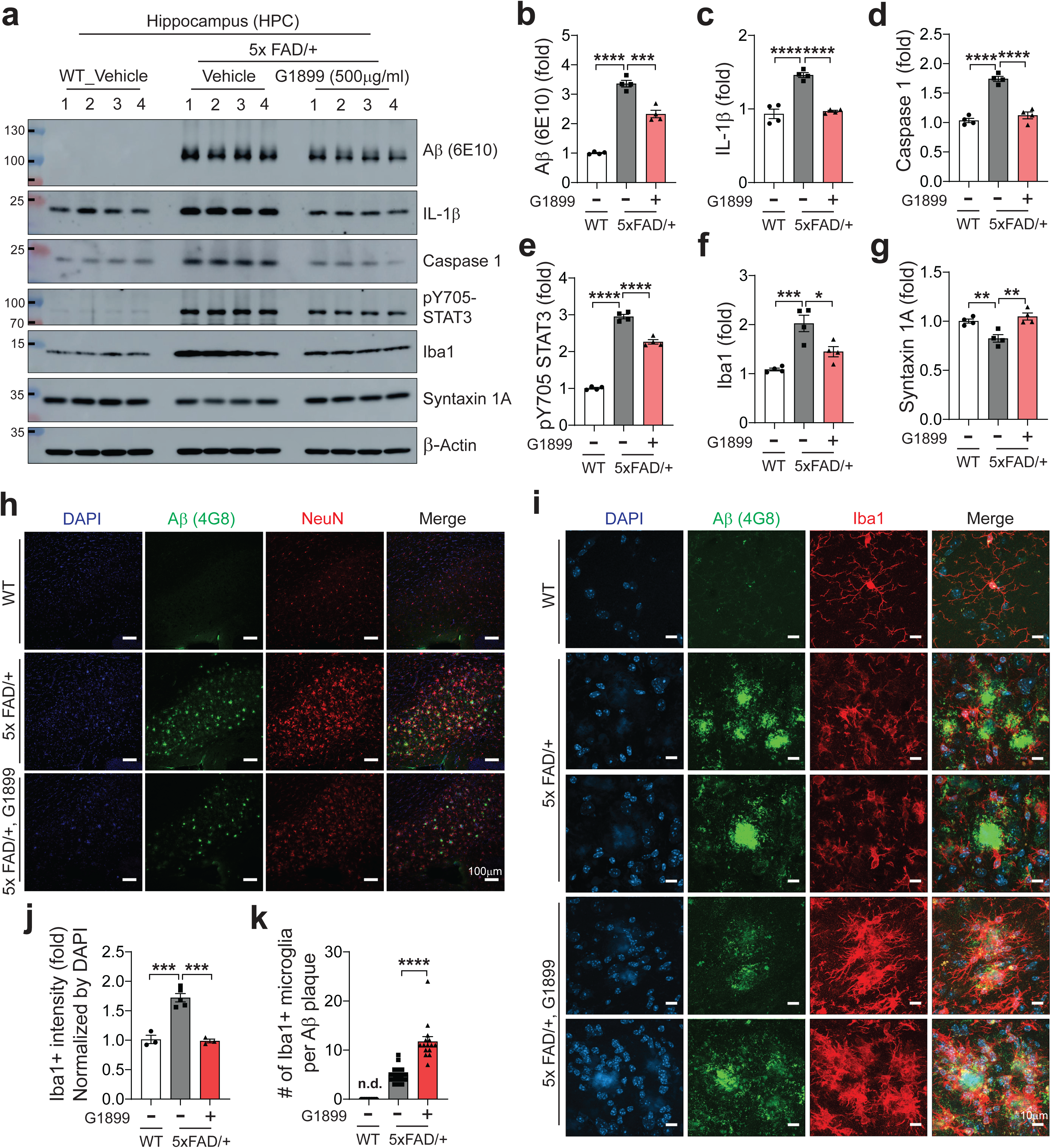
Attenuation of hippocampal neuroinflammation by G1899 in 5xFAD/+ mice. **a** Representative immunoblot images showing expression of Aβ, IL-1β, caspase-1, pY705 STAT3, Iba1, and Syntaxin 1A in the hippocampus of WT, 5xFAD/+, and 5xFAD/+ mice treated with 300 mg/kg G1899. Quantification of relative protein expression of **b** Aβ (6E10), **c** IL-1β, **d** caspase-1, **e** pY705 STAT3, **f** Iba1, and **g** Syntaxin 1A. **h** Representative immunofluorescence images detecting Aβ (4G8) co-stained with Iba1 in the dorsal subiculum of the hippocampus, shown at 10× magnification for global Iba1 intensity (Scale bar, 100 μm) and **i** 63× magnification to visualize microglial recruitment toward Aβ plaques (Scale bar, 10 μm). Quantification of **j** Iba1 immunofluorescence intensity and **k** the number of microglia recruited toward Aβ plaques. Error bars indicate mean ± SEM. Statistical significance is determined by one-way ANOVA followed by Tukey’s multiple comparisons test. *p < 0.05, **p < 0.01, ***p < 0.001, ****p < 0.0001; n.s., not significant; n.d., not detected.

### Beneficial effects of microglia on Aβ-induced neuroinflammation in the context of induced human microglia (iMG)

Building upon our in vivo data demonstrating that G1899 effectively modulates detrimental microglial activation and enhancing phagocytic effect, we sought to validate these findings in human-relevant microglia in the context of Aβ-mediated neurodegenerative condition rather than in a mouse model of AD. To investigate the effects of G1899 in human microglia, we employed human embryonic stem cells (H9 line) and induced their differentiation into microglia. We directed the cells toward a hematopoietic stem cell lineage, followed by microglial differentiation and maturation using commercially available kits from STEMCELL Technologies specifically designed for hematopoietic induction, initiation of microglial differentiation and terminal maturation. We first assessed the efficiency and fidelity of the differentiation process using stage-specific markers **(Fig. 6a-g)**. Upon obtaining fully matured induced human microglia (iMG), we examined both the neuroinflammatory mechanisms induced by Aβo and the potential protective effects of G1899. To simulate a pathological condition, we exposed iMGs to Aβo. Aβo treatment significantly increased the expression of proinflammatory markers, including NLRP3 and IL-1β, whereas this elevation was markedly reduced in iMGs treated with G1899 **(Fig. 6h)**. Consistent with the previously described STAT3-mediated microglial activation pathway in primary murine microglial cultures, Aβo-treated iMGs showed increased level of phosphorylation of STAT3 at Y705 and S727 rather than changes in total form of STAT3, along with elevated level of microglial activation marked by IL-1β, NLRP3, and Iba1. Those elevated protein expressions were effectively reduced by G1899 treatment **(Fig. 6i-n)**. Furthermore, Aβ levels in iMGs treated with Aβo was markedly increased as assessed by Aβ antibodies (6E10 and 4G8). The level of Aβ was significantly reduced in the G1899-treated group, supporting the hypothesis that G1899 promotes Aβo clearance by enhancing phagocytic activity **(Fig. 6o, p)**. In this regard, we sought to validate enhancement of phagocytic activity in G1899 treated group. We firstly observed a dose-dependent increase in the expression of homeostatic microglial marker TMEM119 and the phagocytic marker CD68. These results suggest that G1899 maintains microglia in a beneficial, homeostatic state while enhancing their phagocytic activity, as evidenced by elevated TMEM119 and CD68 levels, thereby supporting a robust defense mechanism against external antigens **(Fig. 6q-s)**. Furthermore, Immunocytochemical analyses further revealed that Aβo-treated iMGs exhibited abnormal morphological changes consistency with reactive microglial states, including amoeboid and rod-like or bushy shapes. However, G1899 treatment reversed these aberrant morphologies, promoting a less reactive, more homeostatic phenotype. Quantification for the cellular area suggested significant morphological differences among the experimental groups **(Fig. 6t, u)**.

**Fig 6.**
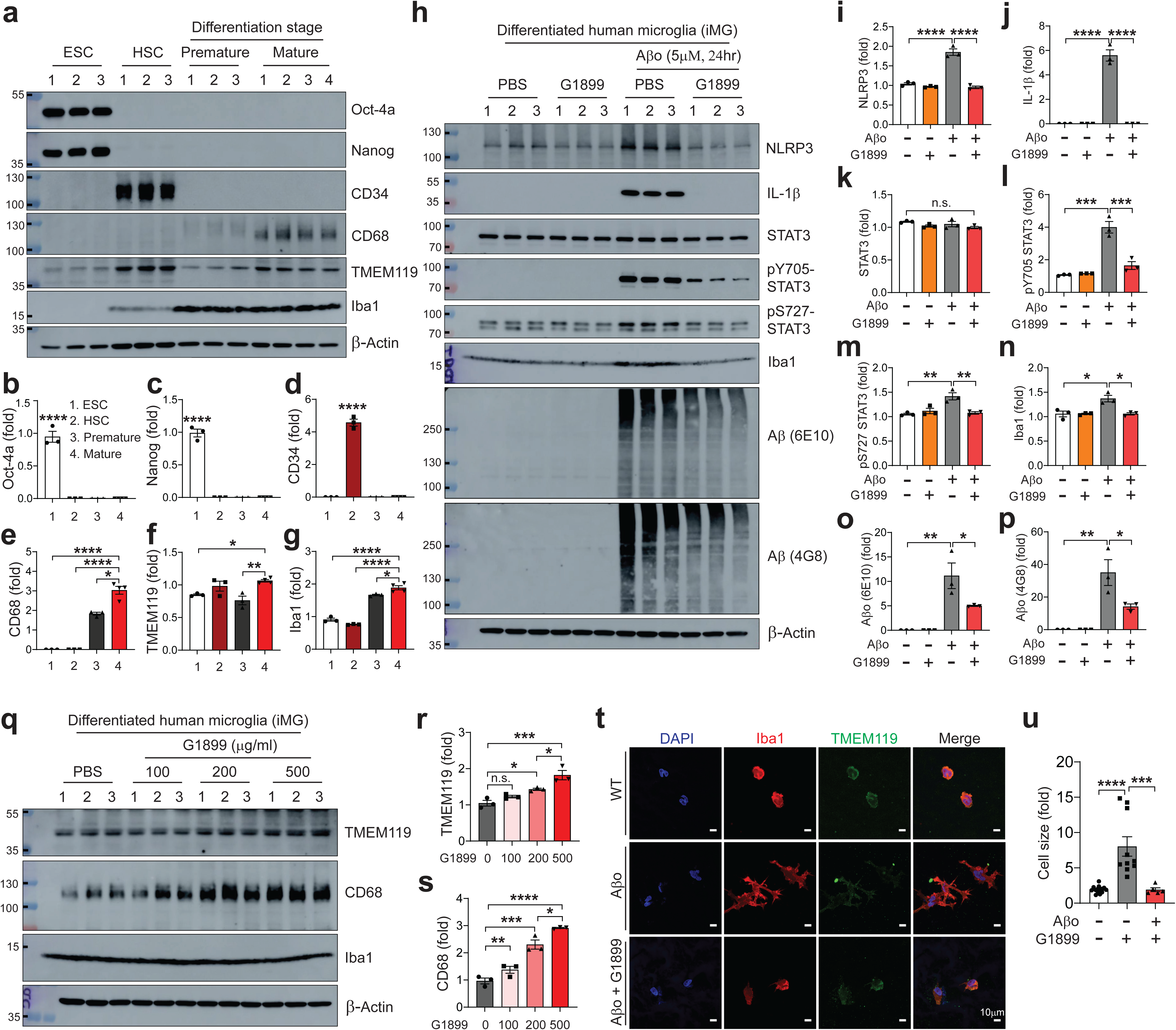
Suppression of inflammatory activation by G1899 in human induced microglia (iMG). **a** Representative immunoblot images showing expression of Oct-4a, Nanog, CD34, CD68, TMEM119, Iba1, and β-actin during stepwise stages of iMG differentiation. Quantification of **b** Oct-4a, **c** Nanog, **d** CD34, **e** CD68, **f** TMEM119, and **g** Iba1 expression across differentiation stages. **h** Representative immunoblot images showing expression of NLRP3, IL-1β, STAT3, pY705 STAT3, pS727 STAT3, Iba1, Aβ (6E10 and 4G8), and β-actin. Quantification of **i** NLRP3, **j** IL-1β, **k** STAT3, **l** pY705 STAT3, **m** pS727 STAT3, **n** Iba1, **o** Aβ (6E10), and **p** Aβ (4G8). **Q** Representative immunoblot images showing expression of TMEM119, CD68, Iba1, and β-actin, with quantification of **r** TMEM119 and **s** CD68. **t** Representative immunofluorescence images of iMG co-stained with Iba1 and TMEM119. Scale bar, 10 μm. **u** Quantification of relative cell size in Fig. 6t. Error bars represent mean ± SEM. Statistical significance was determined by one-way or two-way ANOVA followed by Tukey’s multiple comparisons test. *p < 0.05, **p < 0.01, ***p < 0.001, ****p < 0.0001; n.s., not significant.

## DISCUSSION

Alzheimer’s disease (AD) is a chronic and progressive brain disorder characterized by cognitive impairment, Aβ plaque deposition, and neuroinflammation. Given the limited efficacy of current treatments, identifying alternative therapeutic strategies is critical. American ginseng, a well-known traditional herbal remedy, contains bioactive compounds such as ginsenosides with antioxidant, anti-inflammatory, and neuroprotective properties, making it a promising candidate for AD intervention. Our study systematically evaluated the neuroprotective effects of G1899, an American ginseng extract, in both cellular and animal models, focusing on its potential to modulate neuroinflammation, enhance cognitive function, and mitigate AD-related pathology. American ginseng is a well-established natural supplement with minimal toxicity. Its primary bioactive compounds, ginsenosides, have demonstrated cholinergic system modulation, oxidative stress reduction, and neuroprotection. Unlike existing products, G1899 was extracted using a modified method designed to yield a more stable and enriched profile of bioactive molecules. Based on this, our study placed priority on evaluating G1899’s efficacy in mitigating AD pathology to support its potential as a therapeutic intervention and dietary supplement. Our in vitro experiments demonstrated that G1899 protects neuronal cells from glutamate- and Aβ oligomer (Aβo)-induced toxicity. Since AD pathology involves not only intrinsic neuronal vulnerability but also cytotoxicity mediated by surrounding cells, we extended our investigation beyond neuron-specific protection to evaluate the role of G1899 in microenvironments influenced by non-neuronal cells. In particular, we examined its impact on microglial activation, which is a central process in AD-associated neuroinflammation. G1899 significantly modulated microglial activation, as shown by immunoblot analysis that revealed dose-dependent increases in TMEM119 and CD68 expression. Furthermore, G1899 effectively suppressed proinflammatory signaling pathways triggered by Aβo, including activation of the NLRP3 inflammasome, IL-1β, Caspase-1, and microglial reactivity validated by Iba1. To assess the cognitive-enhancing effects of G1899, we employed both scopolamine (SCP)-induced memory impairment model and the 5xFAD transgenic mouse model of AD. The SCP model provides a robust platform for testing cognitive enhancers and elucidating mechanisms of memory impairment. SCP induces oxidative stress, neuroinflammation, and neurotransmission disruption in key brain regions such as the hippocampus and prefrontal cortex [37–39], making it a standard model for cognitive dysfunction. In this model, behavioral assessments including fear conditioning (FC) demonstrated that G1899 significantly rescued learning and memory deficits as comparable to the positive control, Donepezil. To further examine dose dependency, FC testing revealed a significant rescue effect in the high-dose group (300 mg/kg). Based on these results, we subsequently evaluated G1899 in the 5xFAD mouse, a more chronic and genetically relevant model of AD. Starting at 2 to 3 months of age, mice received G1899 at 30, 100, or 300 mg/kg by oral gavage five times per week for six months followed by behavioral evaluations, including FC and Morris Water Maze (MWM), which revealed that significant improvements in spatial learning and memory in the high-dose group. In the 5xFAD model, long-term administration of G1899 (at high dose of 300 mg/kg) resulted in reduced Aβ plaque burden, suppression of neuroinflammatory markers (NLRP3, IL-1β, Caspase-1, STAT3), and attenuation of microglial activation. To examine microglia-dependent neuroinflammation in a more human-relevant condition, we utilized induced human microglia (iMG) and confirmed that treatment with G1899 restored Aβo-induced microglial activation and inflammatory signaling. Notably, G1899 reduced expression of key inflammation-related signals such as IL-1β and NLRP3. It also modulated the STAT3 signaling pathway, which plays a critical role in microglial inflammatory responses [40] along with NLRP3 inflammasome formation [41, 42]. These findings demonstrate that G1899’s regulatory effects on microglial activation observed in primary cultures are reproducible in iMG systems. Our findings indicate that G1899 contributes to the stabilization of microglial homeostasis while simultaneously enhancing their functional competence under pathological conditions. Specifically, the preservation of TMEM119-positive microglia, together with the enrichment of CD68-expressing phagocytic populations, suggests that G1899 promotes a balanced microglial state that is both homeostatic and capable of efficient clearance of pathological substrates. Importantly, when iMGs were challenged with Aβo, this G1899-maintained population demonstrated a heightened proteolytic capacity, as evidenced by the increased degradation of Aβ species. These results highlight a potential mechanism by which G1899 mitigates amyloid burden and attenuates downstream neuroinflammatory cascades, ultimately supporting the preservation of neuronal integrity in the context of AD. The study further benefits from several strengths, including the use of complementary models spanning neurons, primary microglia, human iMG, and both acute and chronic in vivo paradigms, combined with multimodal readouts that integrated molecular, imaging, and behavioral endpoints. Together, these approaches provided convergent evidence that G1899 exerts consistent neuroprotective actions across systems relevant to AD pathology. Nonetheless, certain limitations should be noted.

While oral administration of G1899 at 300 mg/kg effectively ameliorated behavioral deficits, reduced neuroinflammatory responses, and protected neuronal integrity in mice, the effects were not consistently observed in a dose-dependent manner. This indicates that additional dose– response studies will be required to define the optimal therapeutic window. Given the favorable safety profile of G1899 documented in prior studies [43] and further supported by our findings, it is reasonable to anticipate that administration at higher concentrations may yield more robust and reproducible therapeutic benefits. Furthermore, although the 5xFAD model captures Aβ pathology, it does not recapitulate tau-driven neurodegeneration, which is another critical aspect of AD [44, 45]. Future studies should investigate the efficacy of G1899 in tauopathy models such as P301S [46] or P301L [47] and other neurodegenerative conditions involving cognitive dysfunction and neuroinflammation. In conclusion, our findings provide strong preclinical evidence that G1899 offers neuroprotective and cognitive-enhancing benefits in AD models. By reducing Aβ deposition, suppressing neuroinflammation, and improving cognitive performance, G1899 shows promise as a candidate for both therapeutic and preventive strategies targeting AD-related cognitive decline. Although further studies are required to optimize dosing, clarify mechanisms, and evaluate translational potential in clinical settings, current study underscores the potential of G1899 as a functional supplement supporting brain health and neuroprotection in aging populations.

## Conclusion

In this study, we demonstrated that American ginseng exerts pleiotropic effects that may play an important role in preventing and mitigating age-related neurodegeneration and associated cognitive decline. Given its low toxicity, American ginseng has the potential to be utilized as a daily dietary supplement with beneficial impact. Importantly, through both animal models and human-derived induced microglia, we confirmed that American ginseng helps maintain microglia in a homeostatic state and suppresses NLRP3 inflammasome formation via STAT3 signaling. This mechanism effectively attenuates Aβ-induced neuroinflammation, highlighting its potential as a safe and multifaceted therapeutic strategy for neurodegenerative diseases. Nonetheless, further studies are warranted to determine optimal dosage, assess long-term safety, and validate efficacy in tauopathy and other neurodegenerative conditions, thereby establishing its translational relevance in clinical settings.

## List of abbreviations

Aβ: Amyloid beta
AD: Alzheimer’s disease
SCP: Scopolamine
PSEN1: Presenilin-1
5xFAD: Five Familial Alzheimer’s Disease
APP: amyloid precursor protein
BACE1: β-secretase
NLRP3: NOD-, LRR-, and pyrin domain-containing protein 3
STAT3: Signal Transducer and Activator of Transcription 3
ASC: Apoptosis-related speck-like protein containing a caspase recruitment domain
IL-1β: Interleukin-1 beta
CD68: cluster of differentiation 68
iNOS: inducible nitric oxide synthase
TMEM119: Transmembrane protein 119 (TMEM119)
iMG: Human induced microglia

## Declarations

### Ethics approval and consent to participate

All experimental procedures involving animals were conducted according to the principles of laboratory animal care (NIH publication No. 86-23, revised 1985). All procedures involving animals were approved by and conformed to the guidelines of the Institutional Animal Care Committee of Johns Hopkins University (Protocol Number MO22M174).

### Consent for publication

Not applicable.

### Availability of data and materials

All data generated in this study are included in this published article. Raw datasets during the current study are available from the corresponding author upon reasonable request.

### Author contributions

S.H.K performed all of the experiments in this article and contributed to the experimental designs, interpretation of data and writing the original draft. J.H.L and B.C.H contributed to interpretation of preclinical studies and provided the compound. H.S.K contributed to the resources, supervision, funding acquisition, validation, manuscript revision and project administration.

## Acknowledgements

Not applicable.

---

**Fig S1.**
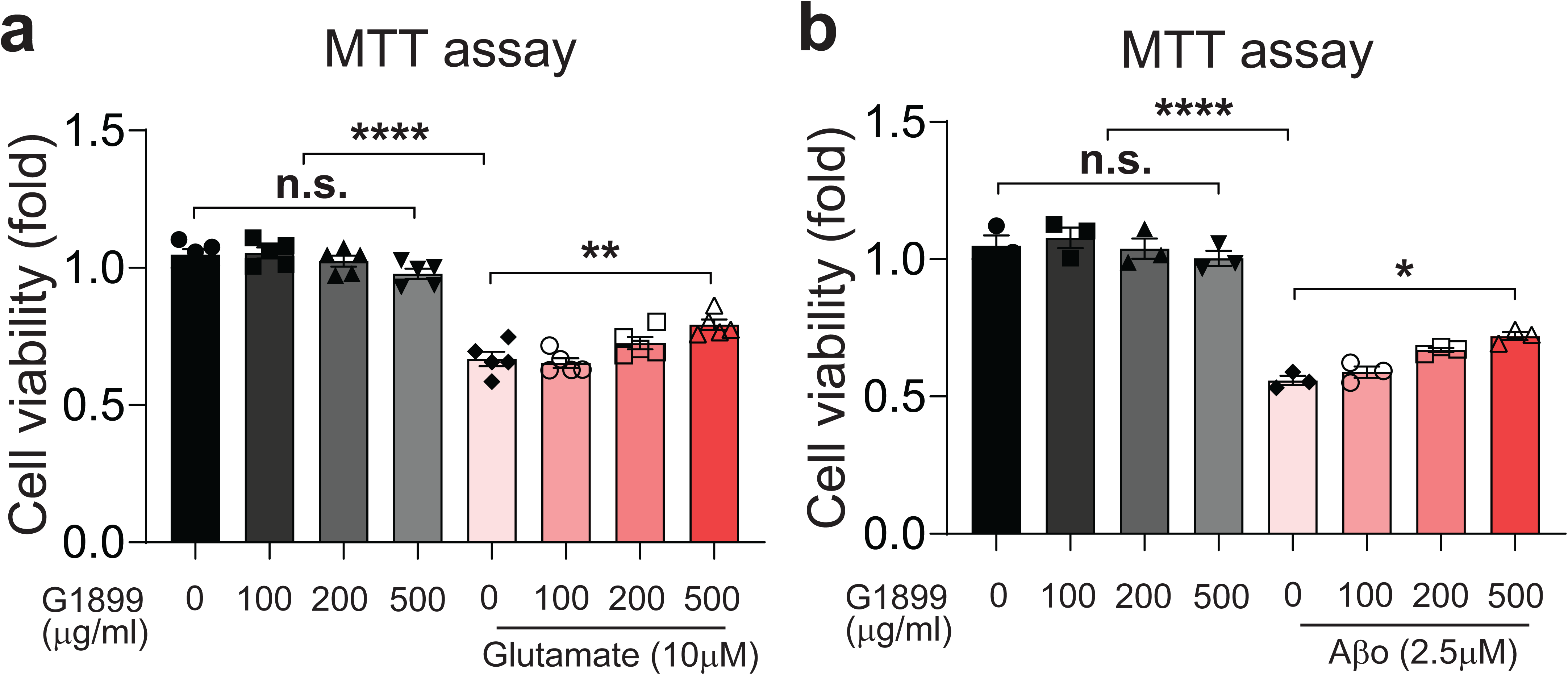
G1899 protects neurons against glutamate- and amyloid β oligomer-induced toxicity. Cell survival following exposure to **a** glutamate (10 μM) and **b** Aβo (2.5 μM) with pretreatment of G1899 at 100, 200, or 500 μg/ml was assessed by MTT assay. Error bars represent mean ± SEM. Statistical significance was determined by two-way ANOVA followed by Tukey’s multiple comparisons test. *p < 0.05, **p < 0.01, ***p < 0.001, ****p < 0.0001; n.s., not significant.

**Fig S2.**
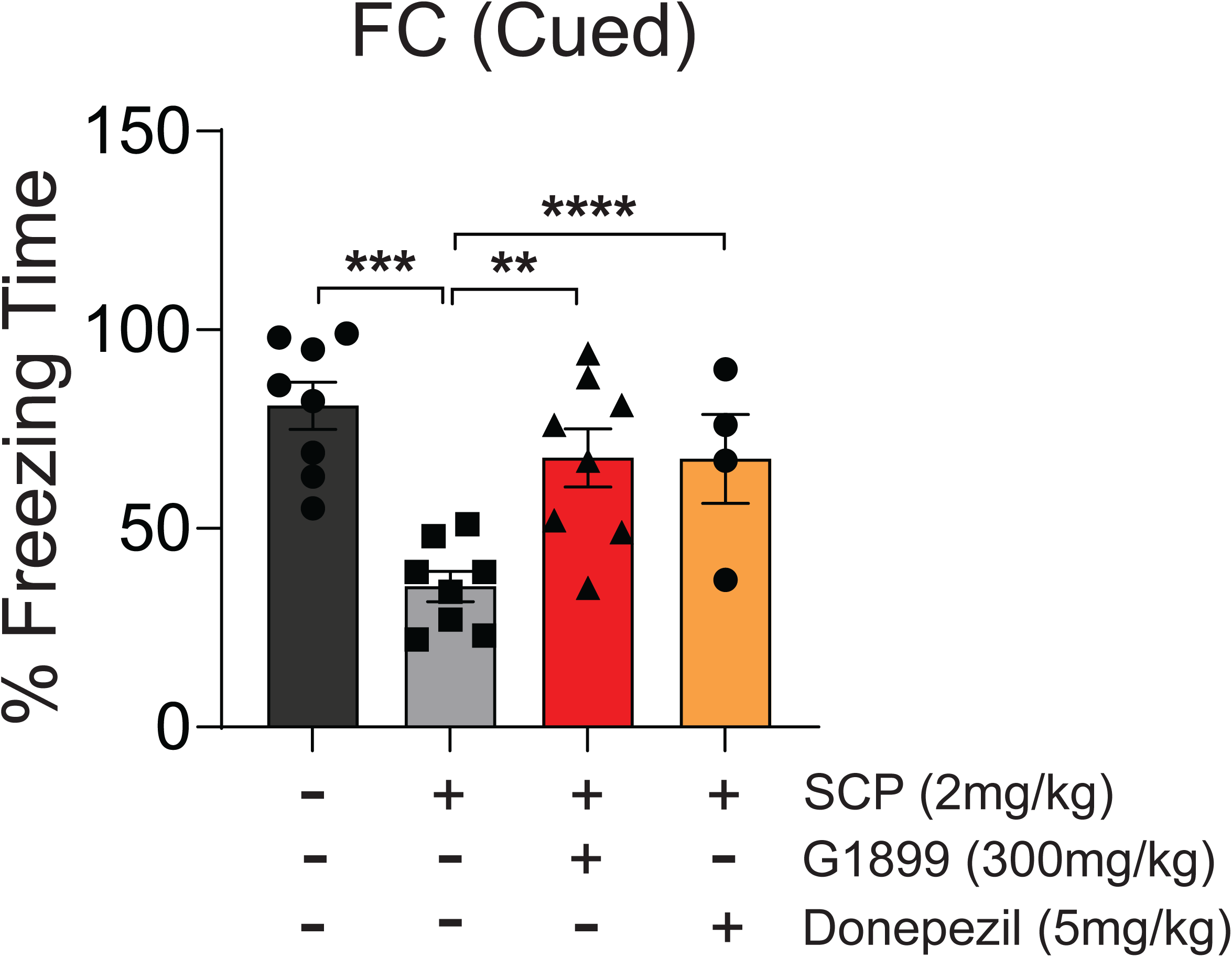
Restoration of cognitive function by G1899 in scopolamine (SCP)-treated mice. Percent of Freezing time was measured by fear conditioning (FC) test to evaluate whether SCP-induced cognitive decline was restored by G1899 treatment, compared with Donepezil (5 mg/kg) as a positive control. Error bars represent mean ± SEM. Statistical significance was determined by one-way ANOVA followed by Tukey’s multiple comparisons test. *p < 0.05, **p < 0.01, ***p < 0.001, ****p < 0.0001; n.s., not significant.

**Fig S3.**
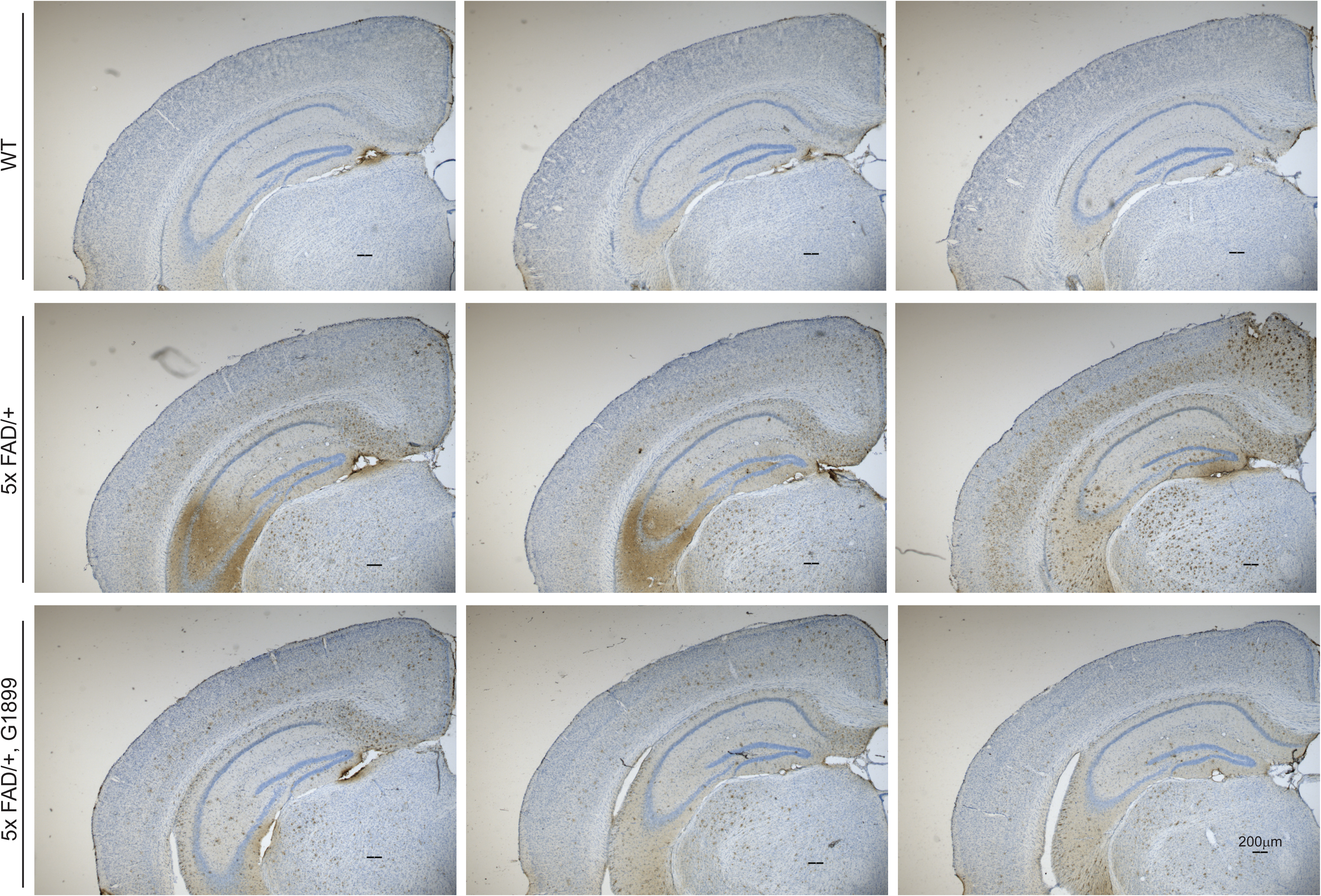
Reduction of amyloid β plaque formation in hippocampal region of 5xFAD/+ mice by G1899. Representative immunohistochemistry images detecting amyloid β (4G8) in the dorsal subiculum of hippocampus of coronal brain sections from WT vehicle, 5xFAD/+ vehicle, and 5xFAD/+ with G1899. Scale bar, 200µm.

**Fig S4.**
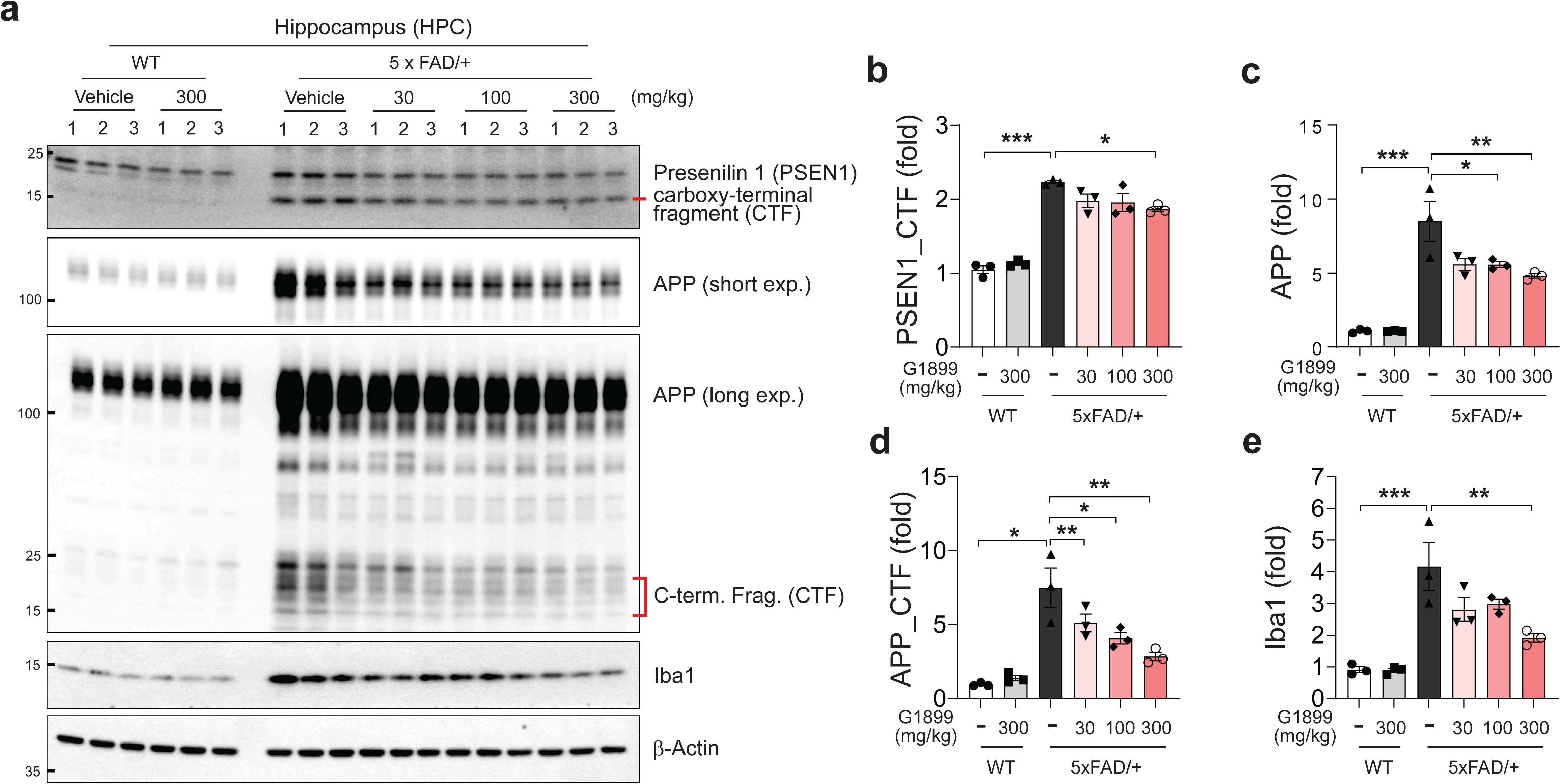
Mitigation of amyloid β-associated pathology in hippocampus of 5xFAD/+ mice by G1899. **a** Representative immunoblot images of hippocampal lysates showing expression of presenilin 1 (PSEN1), APP, and Iba1. Quantification of **b** PSEN1, **c** APP, **d** APP C-terminal fragment (CTF), and **e** Iba1. Error bars represent mean ± SEM. Statistical significance was determined by one-way ANOVA followed by Tukey’s multiple comparisons test. *p < 0.05, **p < 0.01, ***p < 0.001, ****p < 0.0001; n.s., not significant.

